# Minimal requirements and evolution of polar membrane targeting in the land plant SOSEKI protein family

**DOI:** 10.64898/2026.10.30.762996

**Authors:** Andriy Volkov, Evgeniya Pukhovaya, Catherine Albrecht, Dolf Weijers

**Affiliations:** Laboratory of Biochemistry, Wageningen University, Stippeneng 4, 6708WE, Wageningen, the Netherlands; Translational Plant Biology, Utrecht University, Padualaan 8, 3584CH Utrecht, the Netherlands

**Keywords:** Plant cell polarity, SOSEKI, polar protein targeting, evolution

## Abstract

Accurate spatial distribution of polar proteins is essential for cell function, but mechanisms underlying polar protein targeting in plants are poorly understood. Here, we dissect the molecular mechanisms underlying the polar targeting of the SOSEKI1 (SOK1) protein in *Arabidopsis thaliana*. Using extensive mutagenesis and truncation analyses of the polarity domain, we mapped the minimal polarity fragment necessary and sufficient for stable polar targeting to a 25-amino-acid region. This region likely forms an amphipathic helix, and mutations reveal that both its polar and apolar faces are essential for membrane association and polarity. Using phylogenetic analysis, we reconstruct the evolutionary history of the minimal polar targeting region across land plants. Using cross-species expression and protein localization, we infer the ancestral polar SOK localization pattern and show how it remained remarkably conserved, only to diversify upon gene duplication. This work identifies a polar targeting peptide in plants, reveals a mechanism for polar membrane targeting of a plant protein family, and demonstrates both conservation and diversification within this family.

**Summary statement:** Mapping a minimal polar targeting region in SOSEKI reveals a key amphipathic helix. Phylogenetic analysis illuminates an evolutionary path of conservation and diversification of polar targeting.

## Introduction

Cell polarity, the asymmetric distribution of cellular components, is indispensable for organisms. It is crucial for oriented cell division, differentiation, and the development of complex body plans (St. Johnston, 2018). While mechanisms underlying cell polarization and polar protein localization in animals and fungi have been studied in detail (Chiou et al., 2017) (Butler and Wallingford, 2017), much less is known about plant cell polarity (Muroyama and Bergmann, 2019). Plants evolved their multicellularity independently of animals and fungi (Coelho and Weijers, 2026), and animal and fungal cell polarity proteins and regulators are not conserved (Muroyama and Bergmann, 2019)(Ramalho et al., 2021). However, while polarization mechanisms are largely unknown in plants, there is a range of proteins that have been shown to localize polarly within cells. These include integral membrane proteins such as PIN-FORMED auxin transporters(Vanneste and Friml, 2009), and Boron transporters NIP5 and BOR1 (Takano et al., 2010), as well as peripheral membrane proteins such as BASL (Dong et al., 2009), BRX (Breda et al., 2017) and SOSEKI (SOK)(Yoshida et al., 2019)(van Dop et al., 2020). Most polar proteins in plants have close paralogs, and in several cases, different members in each family display different polar localization patterns. This has been best documented for PIN proteins, where the polar localization appears to be encoded by an intracellular domain (Wiśniewska et al., 2006), but the underlying mechanism and the evolutionary path to diversification is unknown.

SOK proteins have properties that make them especially suitable for studying polar targeting mechanisms. SOK proteins are found across land plants, and in all studied species, are polarly localized (van Dop et al., 2020; Yoshida et al., 2019). Importantly, the 5 Arabidopsis SOK proteins each show different polar patterns (**Fig. 1A-E**)(Yoshida et al., 2019), suggesting both conservation of polar targeting in land plants, and specialization of the precise polar domains. SOK proteins tend to localize to the edge between two specific faces rather than to a defined face of the cell (Yoshida et al., 2019). For example, in *Arabidopsis thaliana* root cortex cells, SOK1 localizes only to the apical inner (center-facing) edge, while SOK2 localizes to the basal inner edge. These localization patterns suggest that SOKs integrate cues from multiple body axes (apical-basal and center-periphery axes) for their localization.

**Figure 1:**
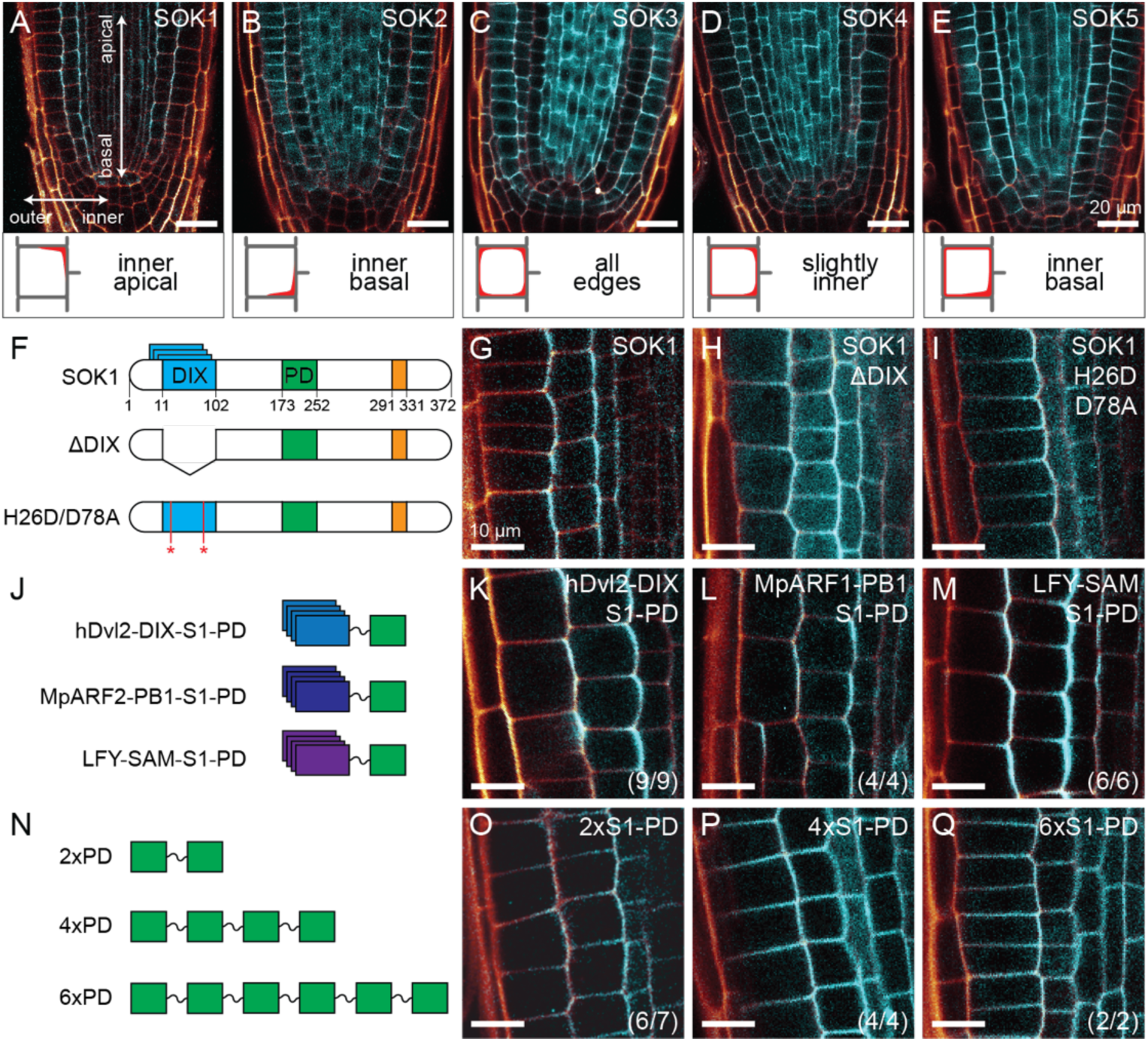
Oligomerization is a requirement for SOSEKI polar membrane localization. (**A**-**E**) Localization of Arabidopsis SOK1 (**A**), SOK2 (**B**), SOK3 (**C**), SOK4 (**D**) and SOK5 (**E**), fused to YFP, and expressed from the *RPS5A* promoter, in Arabidopsis root tips. Coordinates are indicated in (**A**) and interpretations of polar localization patter indicated beneath each root cross-section. (**F**) Cartoons of SOK1 protein variants expressed as RPS5A-driven YFP fusions in (**G**-**I**). (**J**) Cartoons of the fusions between oligomerization domains and the SOK1 polarity domain (PD), expressed as RPS5A-driven mNeonGreen fusions, shown in (**K**-**M**). (**N**) Cartoons of the tandem multimeric PD versions, expressed as RPS5A-driven mNeonGreen fusions (**O**-**Q**). The number of transgenic lines with the pattern shown (n) is shown as a fraction of the total (t) number of lines analyzed (n/t) in each lower right corner. Scale bar for panels (**A**-**E**) is 20 µm and for (**G**-**I**, **K**-**M** and **O**-**Q**) 10 µm.

SOK proteins are peripheral membrane proteins with a conserved domain topology (van Dop et al., 2020). Their defining characteristic is an N-terminal DIX-like domain. This domain is exclusively found in SOK proteins and is highly conserved. The plant DIX domain is structurally and functionally similar to animal DIX domains. Like its animal homologs, plant DIX domains form head-to-tail polymers, binding SOK proteins into higher-order complexes. Following the DIX domain is a long region with no known domains, although there are some conserved sequence motifs. In the middle of the protein is a highly conserved cysteine. Finally, there is a conserved DAxTQT motif just following the DIX domain, as well as an N-terminal Zn finger motif (in some other SOKs, there is a KEY motif instead of the Zn finger).

Previous experiments revealed regions within the SOK1 protein that are critical for polar targeting of SOK1. Firstly, the DIX domain is critical for focusing the protein on an edge. Removing the domain (or interfering with its polymerization) causes the protein to spread out over a larger area of the membrane, albeit retaining membrane localization and polarity (van Dop et al., 2020). Secondly, the conserved cysteine is essential for membrane binding (van Dop et al., 2020; Yoshida et al., 2019). We recently found that this cysteine serves as an acceptor for post-translational S-acylation, through the activity of polar PAT enzymes (Pukhovaya et al., 2026). As a result of this modification, an acyl chain is attached to the cysteine, which anchors the SOK1 protein to the plasma membrane. In experiments, mutating this residue abolished the polarity and membrane localization of SOK1, rendering the protein cytosolic. Lastly, protein truncations and domain swaps identified a fragment of about 80 amino acids surrounding the conserved cysteine to be critical for polarization (Yoshida et al., 2019). Loss of this ‘polarity domain’ (PD) from SOK1 displaces the protein to the cytosol. Moreover, swapping the polarity domain of SOK2 with the equivalent region from SOK1 results in a protein mimicking SOK1 localization, affirming the pivotal role of the polarity domain in selecting the appropriate edge. In summary, previous data suggests that at a minimum, three components are needed to polarize SOKs: DIX-mediated polymerization, the activity of the polarity domain, and S-acylation of a cysteine within the polarity domain.

In this study, we explore the minimal mechanisms guiding polar SOK localization to the membrane. We identify a minimal region within the Arabidopsis SOK1 PD that encodes an amphipathic helix, whose amphipathicity is critical for membrane targeting. We next explore the evolutionary trajectory that this PD followed in the SOK family, and show that most SOK clades retain an ancestral polar localization pattern. This ancestral pattern diversified only after gene duplications occurred in flowering plants. Our work therefore identifies a minimal principle for polar membrane association of SOK proteins and reveals that an ancient, universal polarization system controls the localization of SOK proteins in all plants.

## Results

### Oligomerization of the polarity domain is required for focused edge localization

Previous results demonstrated that Arabidopsis SOK1 and SOK5 proteins rely on their DIX domain for focused localization to the edge, and that the native SOK1 DIX domain could be substituted by a human Dvl2 DIX domain (van Dop et al., 2020). Still, it remains unclear whether only the oligomerization conferred by the DIX domain causes this effect, or whether the DIX domain additionally encodes information pertinent to polar targeting. We first confirmed loss of edge-polarized SOK1 localization in full-length SOK1 (**Fig. 1F,G**) lacking its DIX domain (**Fig. 1H**) or carrying oligomerization mutations (**Fig. 1I**). While these mutants still localize polarly to the membrane, they were more diffuse and less concentrated on the edge compared to the wild-type control. We next replaced the DIX domain of SOK1 with unrelated oligomerization domains (**Fig. 1J**): SAM-SOK1 (carrying a SAM domain from the *Arabidopsis thaliana* LFY protein)(Sayou et al., 2016), and PB1-SOK1 (carrying a PB1 domain from the *Marchantia polymorpha* ARF1 protein)(Kato et al., 2020), each fused to a fluorescent protein. Both oligomerization chimeras localized polarly (**Fig. 1L,M**) and exactly as the hDvl2-DIX swap (**Fig. 1K**). This observation indicates that oligomerization of the polarity domain is sufficient for edge localization. The DIX domain does not encode any positional information and is only responsible for focusing the protein on the edge, but not for polarity itself. Instead, this polarity must be contained in the polarity domain of SOK1.

Oligomerization allows clustering of polarity domains, thus increasing the avidity of the SOK complex for the membrane. To test if increasing avidity up to 6 times that of the monomeric wild-type protein would allow for focused polar edge localization, we expressed (non-oligomerizing) tandem fusions of the SOK1 polarity domain. Specifically, we expressed three constructs, solely consisting of two (dimer), four (tetramer), or six (hexamer) tandem fused copies of the SOK1 polarity domain (2xS1-PD, 4xS1-PD, 6xS1-PD; **Fig. 1N**). All three constructs localized polarly to the membrane but diffusely (**Fig. 1O-Q**), similar to the non-oligomerizing SOK1-H29D/D78A mutant (**Fig. 1I**). This demonstrates that significantly more than six units are needed for edge localization, and/or oligomerization serves to not only increase the effective number of polarity domains but (also) contributes to the localization pattern in another way.

### Mapping of a minimal polarity domain

Having established that polar targeting of SOK1 is encoded in its polarity domain, we further delineated the minimal requirements for polar targeting within this domain. Previously, the boundaries of the polarity domain of SOK1 were based on a series of deletion mutants and a domain swap between SOK1 and SOK2 (Yoshida et al., 2019). However, it remains unclear whether the entire 80-amino-acid fragment is required for polar localization. We substituted parts of the SOK1 polarity domain with flexible glycine-serine linkers in blocks of 10 amino acids, yielding eight mutants covering the entire polarity domain except the essential CG motif (**Fig. 2A**), and expressed each as a fusion to the hDvl2-DIX domain and a fluorescent protein (**Fig. 2B-J**). We scored the localization as membrane-associated if at least some signal was concentrated at the polar membrane domain, even if the cytosolic background signal was higher than in the controls. Note that the use of the FastGreen marker for transformation, which drives strong green fluorescence in seeds (Shimada et al., 2010), causes persistent accumulation of large fluorescent particles in some roots that can not be fully spectrally separated from the mNeonGreen used for protein imaging (**Fig. 2,3,5**). Hence, we disregard such large cytosolic structures and focus only on membrane-associated and homogeneous cytosolic fluorescence. Even if there was variation in these patterns among individual transgenics, the interpretation of the localization pattern was supported by all lines tested. Mutants replacing the first 40 amino acids hardly differed from the non-mutated control (scans 1-4; **Fig. 2C-F**). However, scans 5 and 6 dramatically decreased membrane association and localized to the cytosol instead (**Fig. 2G,H**). Of the remaining mutants, scan 7 showed a mixed cytosolic and polar membrane localization, with a weak membrane localization and a significant cytosolic fraction (**Fig. 2I**), while scan 8 localized polarly (**Fig. 2J**). Interestingly, all the proteins investigated in this experiment were either membrane-bound and polar, or cytosolic, and we never found membrane-localized but apolar mutants, or mutants whose polar domain was modified. In summary, this experiment demonstrates that about 30 amino acid residues contained in scans 5-7 are necessary for polar edge localization, while the others are dispensable.

**Figure 2:**
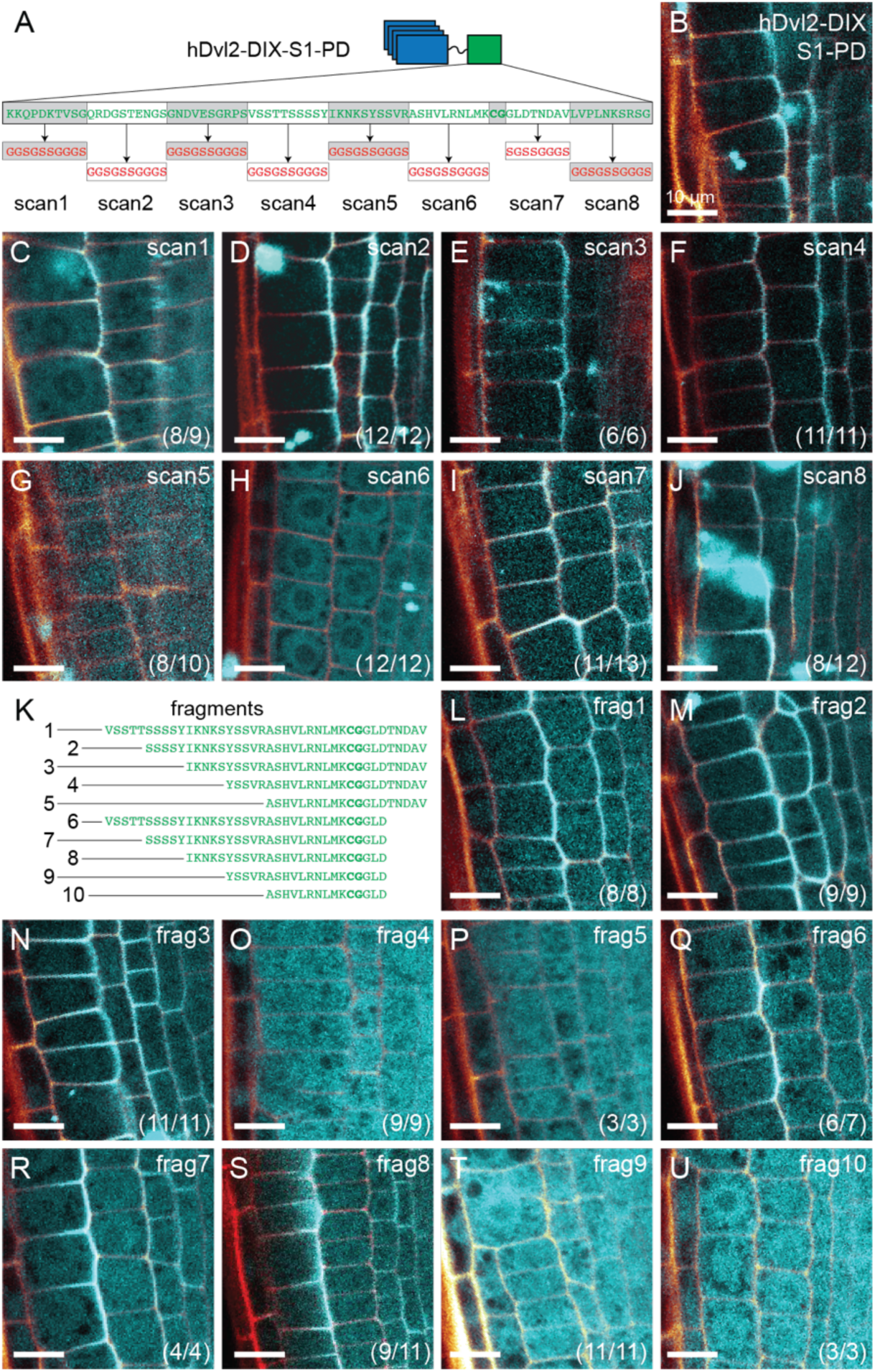
Identification of a minimal region for SOK1 polar membrane targeting. (**A**,**K**) Cartoon showing the substitution scans (**A**) and truncations (**K**) generated within the SOK1 PD, expressed as *RPS5A*-driven fusions to hDvl2-DIX and mNeonGreen and localization of the substitution scans (**B**-**J**) localization and truncations (**L**-**U**) in Arabidopsis root tips. The number of transgenic lines with the pattern shown (n) is shown as a fraction of the total (t) number of lines analyzed (n/t) in each lower right corner. Scale bar for panels (**B**-**J** and **L-U**) is 10 µm.

**Figure 3:**
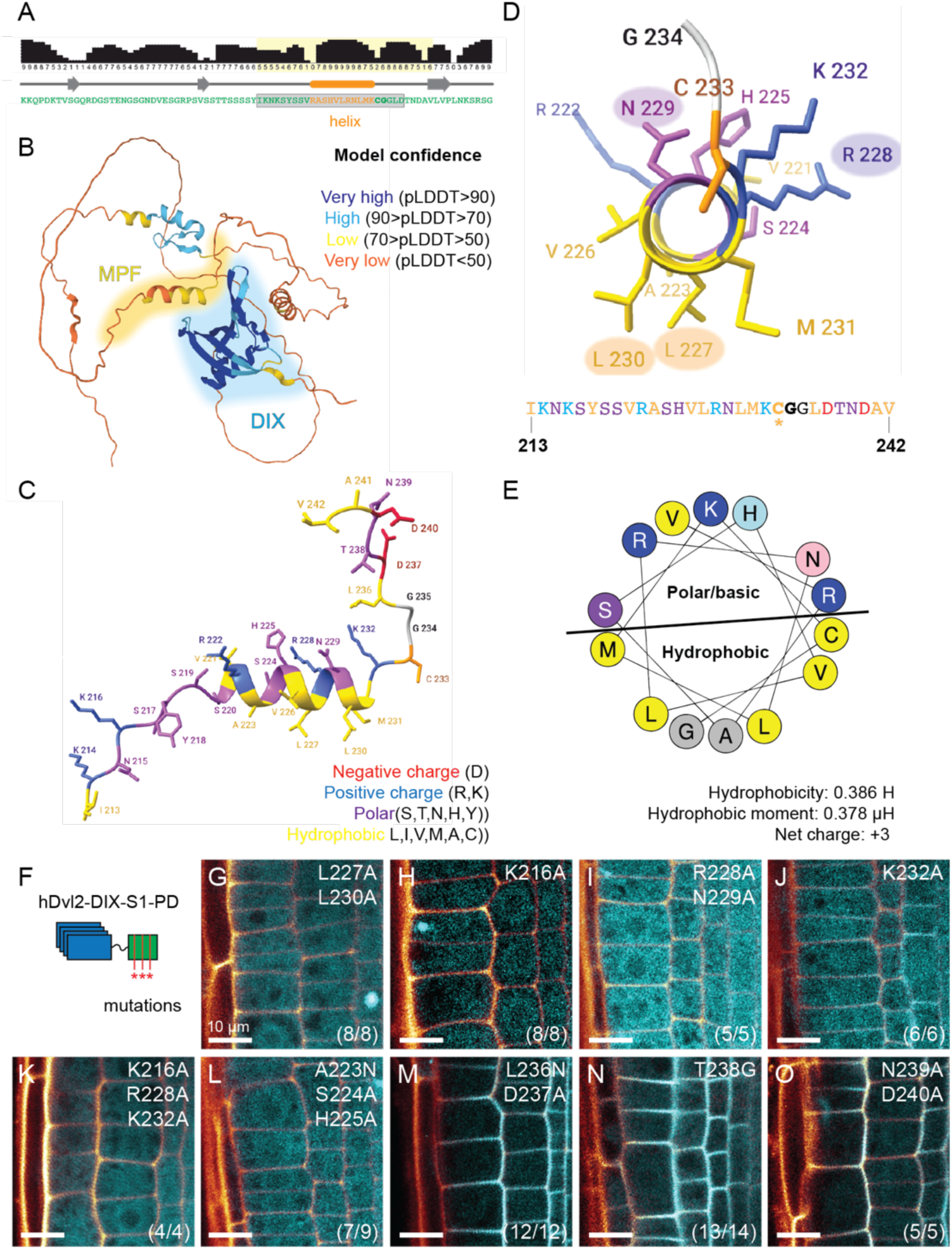
An amphipathic helix drives SOK1 polar membrane targeting. (**A**) Secondary structure prediction (JPRED) for the SOK1 PD region, showing an alpha helix (orange) within the CPF region (boxed). (**B**) AlphaFold2 prediction of the SOK1 structure, showing high-confidence DIX domain structure and a low-confidence helix within the MPF region (both highlighted). (**C**) Detail of the MPF region, showing the predicted helix with side chains, colored by physicochemical properties. (**D**) Alpha helix viewed from different angle, with its sequence underneath, and residues color-coded by properties. (**E**) Helical wheel projection of the helix showing polar/basic and hydrophobic faces. Calculated amphipathic helix properties are indicated. (**F**) Cartoon showing the mutations generated within the SOK1 PD, expressed as *RPS5A*-driven fusions to hDvl2-DIX and mNeonGreen, and localization of mutated fragments (**G**-**O**) in Arabidopsis root tips. The number of transgenic lines with the pattern shown (n) is shown as a fraction of the total (t) number of lines analyzed (n/t) in each lower right corner. Scale bar for panels (**G**-**O**) is 10 µm.

To investigate which region within the polarity domain is sufficient for polar targeting, we generated N- and C-terminally truncated versions of the SOK1 polarity domain (**Fig. 2K**), and likewise expressed each as a fusion to hDvl2-DIX and a fluorescent protein. Each fragment again conferred either polar membrane localization, cytosolic localization, or a mix of both (**Fig. 2L-U**). The minimal fragment required for consistent polarization is encoded by 25 amino acids, namely residues 213-237 of the SOK1 protein (Fragment 8; **Fig. 2S**). A C-terminally extended Fragment 3 did however, appear to subtly enhance membrane localization relative to Fragment 8 (**Fig. 2N**), suggesting that there is a quantitative contribution of the residues C-terminal to Fragment 8. Nonetheless, given that the 25 amino acids are sufficient for polar targeting, we consider it the minimal polarity fragment (MPF).

### A predicted amphipathic helix in the SOK1-MPF mediates polar targeting

Having identified the minimal SOK1 fragment required for stable polar targeting, we wondered how such a short fragment confers membrane localization and polarity. We analyzed sequence features and used prediction tools to explore the nature of this region, and used the extended motif (Fragment 3; **Fig. 2K,N**). Both JPred and AlphaFold predict an α-helix within this fragment (**Fig. 3A,B**). We also noticed the abundance of charged amino acids (**Fig. 3C**): five out of 30 residues are positively charged (R and K) and only two are negatively charged (D). Closely examining the predicted AlphaFold structure of this polarity fragment showed that most of the charged amino acids are on one face of the α-helix, while the other face is abundant in hydrophobic amino acids, including two highly hydrophobic leucines (**Fig. 3C,D**). A helical wheel projection showed clear amphipathicity of the helix through grouping of positive residues on one face of the helix and hydrophobic residues on the other (**Fig. 3E**). Such amphipathic α-helices (AHs) are well-known to be involved in the membrane binding of various proteins (Lee et al., 2026). These proteins typically rely on positive charges (conferred by R and K residues) on one face of an AH for recruitment to the negatively charged phospholipid head groups (Giménez-Andrés et al., 2018). The hydrophobic face of the helix ensures association within the lipid bilayer through long-chain aliphatic residues (frequently L). This configuration of amino acids confers high hydrophobic moments to the helix. In our case, the hydrophobic moment is as high as 0.378 µH, which is typical for membrane-associated helices (Eisenberg et al., 1982). The molecular dynamics tool PMIpred predicts strong binding of the AH to negatively charged membranes with a high free energy of membrane binding (**Fig. S1**). These simulations of binding kinetics suggest that the SOK1 MPF acts to bind membranes.

We next directly tested the role of amphipathicity within this predicted AH by targeted mutations in the context of Fragment 1 (**Fig. 2K,L**; **Fig. 3F**). Mutating the two leucines on the hydrophobic face of the helix (L227A/L230A) completely abolished membrane association (**Fig. 3G**). Similarly, to manipulate the hydrophilic face of the AH, we mutated various charged and hydrophilic residues. Mutants with altered positive charges (K216A; R228A/N229A; K232A; K216A/R228A/K232A) were similarly cytosolic and failed to localize to the membrane (**Fig. 3H-K**). The K232A mutant showed an intermediate effect by localizing both polarly to the membrane and to the cytosol (**Fig. 3J**), suggesting it is less critical than K216. Other residues within the helix are also essential, as demonstrated by the cytosolic mutant A223N/S224A/H225A (**Fig. 3L**), while mutations C-terminal of the cysteine (and therefore outside of the amphipathic helix) did not affect localization (L236N/D237A; T238G; N239A/D240A; **Fig. 3M-O**). Together, these findings show that hydrophobicity and positive charge on either face of the AH, respectively, are essential for SOK1 polar membrane association.

### The minimal polar targeting region shares conserved features in SOSEKI proteins

The five Arabidopsis SOSEKI proteins differ in subcellular targeting (Yoshida et al., 2019) (**Fig. 1A-E**), which indicates that the targeting mechanisms and sequences must have diverged. Previously, we reconstructed the evolutionary history of the SOSEKI protein family (van Dop et al., 2020) (Ramalho et al., 2021). However, the density of sampling in some clades was too limited to infer patterns of evolution at the scale needed to interpret evolution of the polarity domain. We therefore developed a more extensive phylogenetic tree (**Fig. 4A,B**) using the most recent available genomic data. This phylogeny includes 1086 SOK sequences from 199 plant species. Relative to previous studies (Ramalho et al., 2021)(van Dop et al., 2020), this phylogeny includes a large number of angiosperm sequences, as well as additional species of lycophytes and bryophytes. Additionally, by reconciling the SOK gene tree with a known species tree, we resolved recent and ancient evolutionary events. Our data are consistent with previous analyses (van Dop et al., 2020), but add detail about the origin of SOK orthologs. We infer that a single ancestral SOK (AncSOK) in early embryophytes (land plants) founded the single bryophyte SOK ortholog group, BryoSOK. These BryoSOKs form a sister clade to all tracheophyte (vascular plant) SOKs, which independently evolved from AncSOK. The single AncSOK underwent a series of duplications and speciations in the tracheophyte lineage. From these events, several ortholog groups arose, of which the lycophyte-specific LycoSOK is the earliest-diverging group. All euphyllophyte SOKs form a single EuSOK ortholog group. This group diversified into several subgroups. At the base of euphyllophytes, the ancestral EuSOK duplicated, forming the two major ortholog groups EuSOK-A (containing AtSOK1) and EuSOK-B (containing AtSOK2-5). Both groups are broadly found across all euphyllophyte lineages. We further detected a recent diversification of EuSOK-B genes into three subgroups. A series of duplications at the base of spermatophytes (seed plants) and angiosperms (flowering plants) formed three angiosperm-specific ortholog groups, namely EuSOK-B2 (including AtSOK2), EuSOK-B3 (including AtSOK3), and EuSOK-B4 (including AtSOK4/5). All the major angiosperm groups (including basal angiosperms, monocots, and dicots) retained SOKs from each of these three groups, suggesting that each group is relevant for flowering plants.

**Figure 4:**
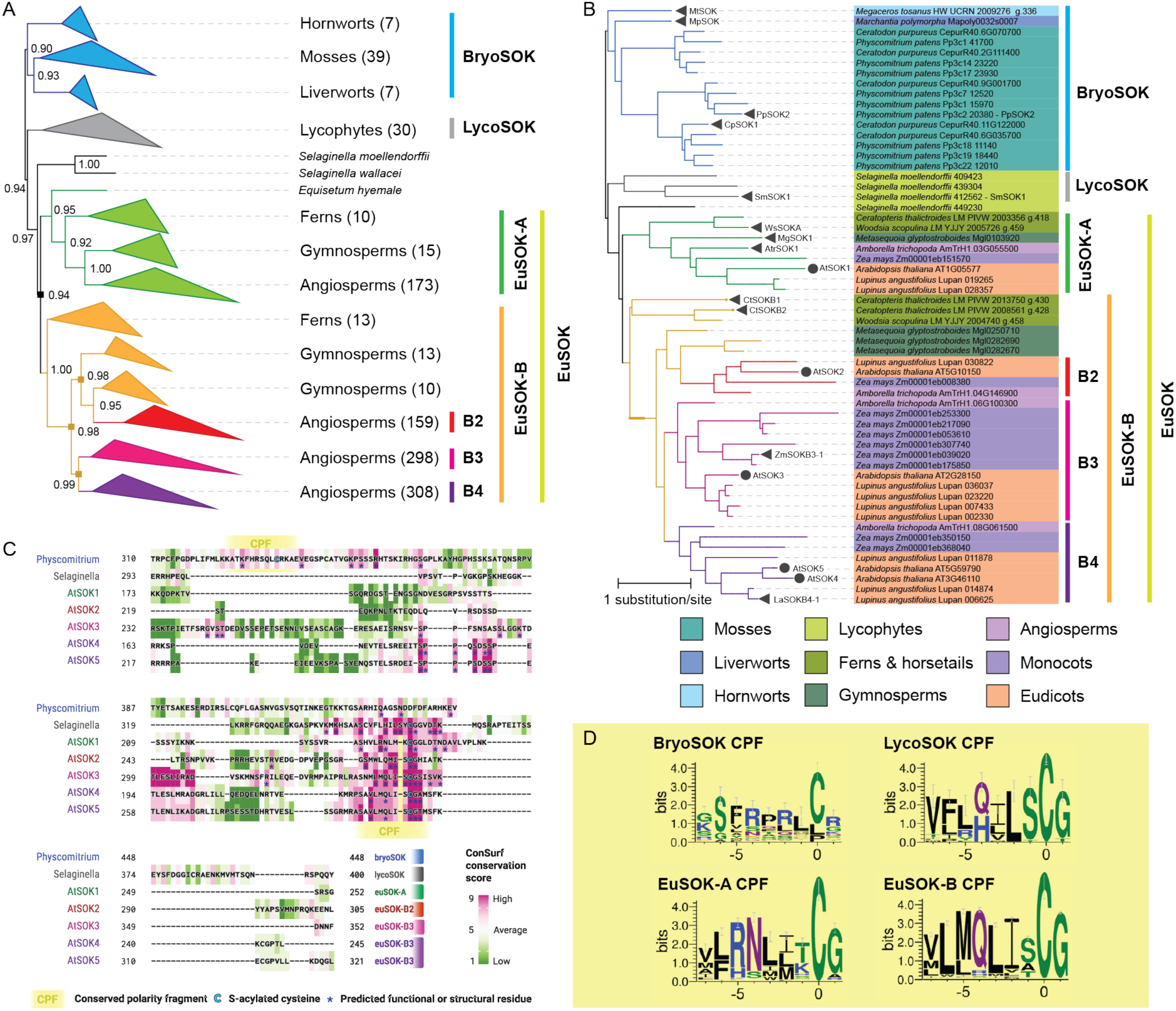
SOK phylogeny identifies conserved motifs within the polarity domain. (**A**) Simplified phylogenetic tree for the SOSEKI protein family, identifying deeply conserved clades in Bryophytes (BryoSOK), Lycophytes (LycoSOK), Euphyllophytes (EuSOK), subclasses within Euphyllophytes (EuSOK-A, EuSOK-B) and within Angiosperms (EuSOK-B2, -B3 and -B4). Bootstrap values are indicated at each node. (**B**) Abbreviated phylogenetic tree highlighting all relevant clades, and species are color-coded by phylogenetic distribution (see legend at bottom). Arabidopsis (AtSOK) sequences are marked, and sequences selected for functional analysis are marked with triangles. The scale bar shows branch length relative to the substitution rate. (**C**) Alignment of the Polarity Domain in all 5 Arabidopsis SOK proteins, as well as individual Selaginella and Physcomitrium shown (unaligned) SOK proteins (thus reflecting all clades), highlighting conserved motifs and residues through ConSurf color coding. (**D**) Sequence logos for the identified conserved motifs in each SOK subclasss.

Since different SOK groups, even the more recently diverged ones, have proteins with different localization patterns, we aimed to identify group-specific differences in the polarity domains of SOKs from the groups. We analyzed the regions in BryoSOKs, LycoSOKs, EuSOK-As, and EuSOK-Bs that were in equivalent positions to the 80-amino-acid broader polarity domain of SOK1. Subsequently, we estimated the evolutionary conservation of residues within these domains using ConSurf. Within the polarity domain, we found a conserved region common to all SOK proteins, including the acylated cysteine (**Fig. 4C**). We refer to this region as the Conserved Polarity Fragment (CPF). Except for BryoSOKs, the consensus sequence of this region is always VLxxLxxCG (**Fig. 4D**). ConSurf also predicted functional importance for this CPF in all SOK groups, supporting its critical role in SOK function. Detailed motif analysis revealed differences in the CPF sequence between the SOK groups **(Fig. 4D**). In LycoSOKs and EuSOK-Bs, the CPF is abundant with hydrophobic residues (such as leucine, phenylalanine and methionine). In contrast, the CPF of EuSOK-As invariably contains a positively charged residue, such as arginine, which is likely a derived form. Interestingly, this arginine is indispensable for polarization of AtSOK1 (**Fig. 3I,K**). The CPF of BryoSOKs appears less conserved, but nonetheless often consists of hydrophobic and positive residues. Unlike all other SOKs, the cysteine in BryoSOKs is not usually followed by a glycine. In summary, detailed phylogenetic and motif analysis identified both conserved and divergent features among SOSEKI protein groups.

### Broad conservation of ancestral SOK polarity

Having identified a minimal polar targeting fragment in Arabidopsis SOK1, and having mapped the evolution of this region, we next asked how its function evolved. We selected 13 SOK polarity domains from 12 species covering phylogenetic clades from BryoSOKs to EuSOKs (**Fig. 4B**). Each was expressed as a fusion to hDvl2 DIX and mNeonGreen, and we analyzed localization in Arabidopsis roots (**Fig. 5**). To test if the polar patterns of SOK proteins are preserved within their PD region beyond AtSOK1, we also included the AtSOK3 PD in this analysis. We found this PD to recapitulate the all-edges polar pattern of full-length AtSOK3 (**Fig. 5**). In the set of orthologous PD fragments, we found that, except for one of the fern EuSOK-B sequences (CtSOKB1) and an angiosperm euSOK-B4-type sequence (LaSOKB4-1), which were both (mostly) cytosolic, all proteins polarly localized to the membrane of epidermis and cortex cells. All but two localized to the inner membrane, similar to AtSOK1 (**Fig. 1A**). The Marchantia SOK domain (MpSOK), localized to the outer basal membrane domain. The *Zea mays* EuSOK-B3-type protein localized throughout the membrane and was more concentrated at cell edges (ZmSOKB3-1), exactly like AtSOK3 (**Fig. 1C**). The EuSOKB4-type sequence LaSOKB4-1showed some weak membrane association and behaved similarly to AtSOK4, which is distinctly less polar than other SOK proteins in Arabidopsis (**Fig. 1A-E**)

**Figure 5:**
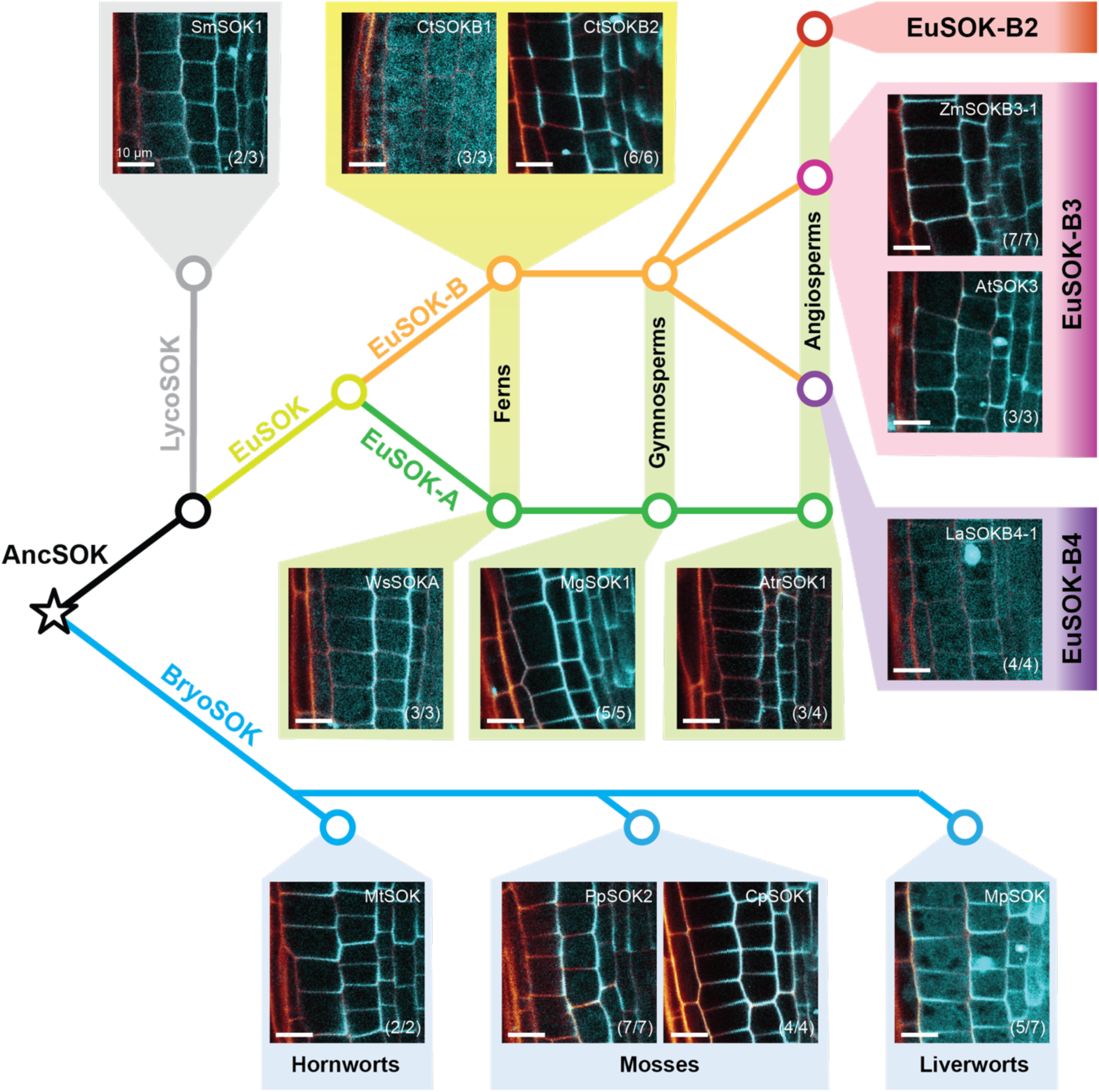
Evolution of polar targeting through the SOK polarity domain. Schematic phylogeny of the SOK family in land plants, with the ancestral proteins (AncSOK, EuSOK) indicated at the relevant nodes and extant SOK classes marked on branches. Microscopy images show the fluorescence of chimaeric proteins carrying hDvl2-DIX fused to the PD domain of 12 land plant SOK proteins (see legend Figure 4 for identity and abbreviations), fused to mNeonGreen and expressed from the RPS5A promoter in Arabidopsis root tips. The number of transgenic lines with the pattern shown (n) is shown as a fraction of the total (t) number of lines analyzed (n/t) in each lower right corner. Scale bar for all microscopy panels is 10 µm.

In summary, this data shows that most tested SOK polarity domains confer the same polar inner membrane localization in Arabidopsis roots, likely reflecting an ancestral character state. While there may be variation in all clades beyond the sequences we experimentally tested, we find that functional diversification within the polarity domain occurred within the euSOK-B clade, likely giving rise to the diverse SOK patterns observed in Arabidopsis.

### Diversification of SOK polarity through the polarity domain

Our comparative analysis demonstrated that the ancestral SOK localization corresponds to the inner lateral domain in Arabidopsis roots, and that later diversification in the EuSOK-B clade generated novel polar localizations. However, MpSOK is a remarkable exception: It localizes to the exact opposite domain when expressed in Arabidopsis roots (**Fig. 5**), compared to all other non-EuSOK-B domains. This suggests that the MpSOK protein has undergone modifications that changed its polar membrane targeting. We have recently shown that MpSOK polar targeting depends on the conserved Cysteine in the polarity domain (Pukhovaya et al., 2026), similar to AtSOK1 and AtSOK5 (van Dop et al., 2020). It is therefore an open question whether the opposite localization of MpSOK in Arabidopsis roots reflects a different wiring of the cellular polarity system, or just altered behavior when expressed in Arabidopsis. To address this question, we performed the reciprocal experiment and expressed AtSOK1 in Marchantia thallus, driven by the EF promoter. In this tissue, native MpSOK polarizes towards the meristematic notch in a supra-cellular, tissue-wide pattern (van Dop et al., 2020) (**Fig. 6A**). Strikingly, within the same cells, AtSOK1 polarized in the opposite pattern, away from the meristematic notch (**Fig. 6B**). Thus, the opposite polarity between AtSOK1 and MpSOK is conserved in the two species, suggesting that the entire cell polarity system that guides SOK polarization is rewired in Marchantia.

**Figure 6:**
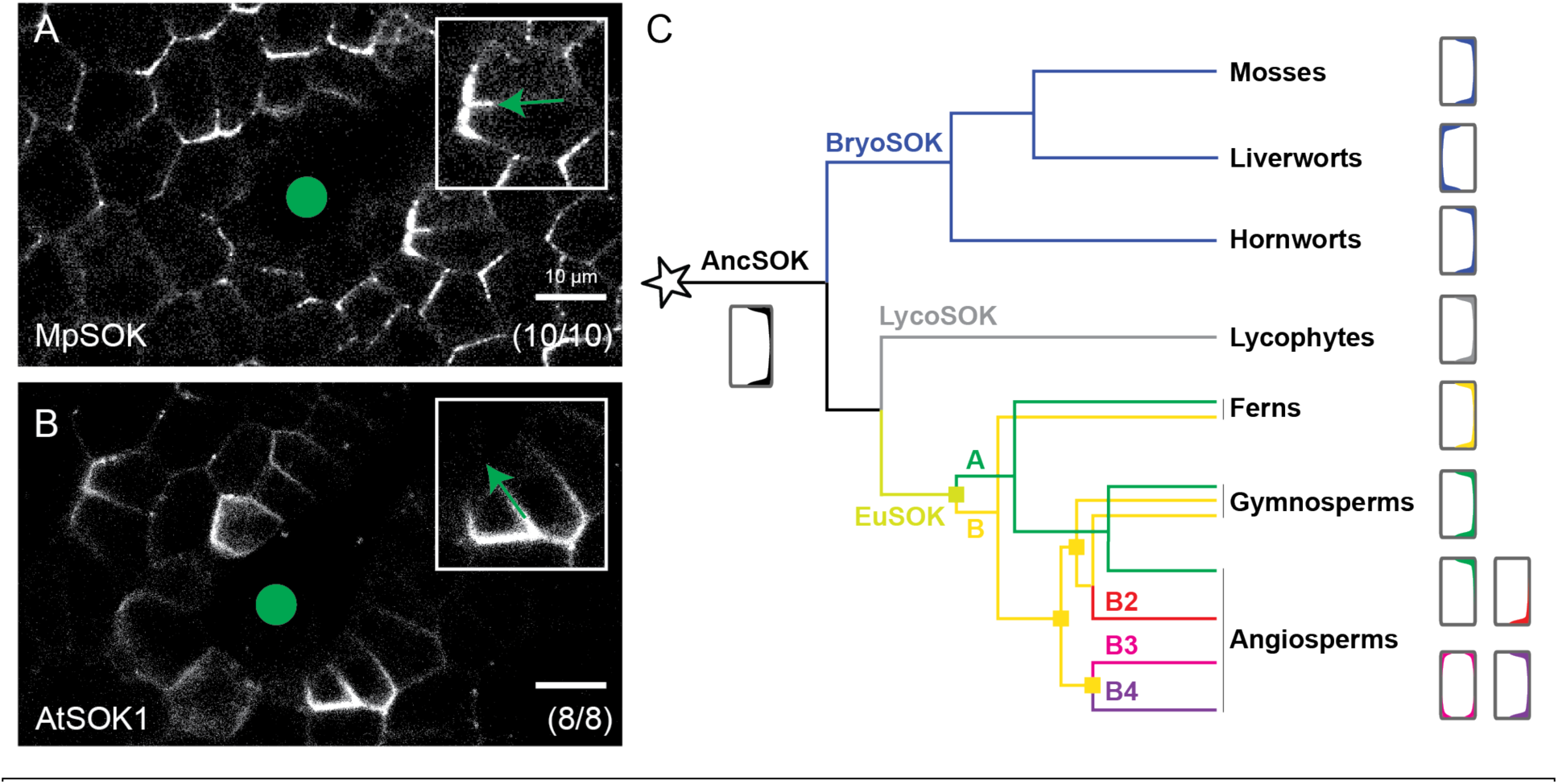
Divergence and evolution of SOK polar targeting. (**A**,**B**) Localization of MpSOK (**A**) and AtSOK1 (**B**), each expressed as a fusion to mCitrine from the *EF* promoter in Marchantia thallus, shows coordinated polarization towards (MpSOK) and away from (AtSOK1) the meristematic notch (green dot). Insets show cells in equivalent position with the direction of the notch marked by an arrow. (**C**) Reconstruction of the evolution of SOK polarity in land plants. The tree is a combined protein sequence tree and species tree, and shows the trajectory of speciations (regular nodes) and duplications (squares at nodes). Inferred ancestral SOK polarity (AncSOK), as well as extant polarity patterns (right) are indicated. Scale bar in (**A**,**B**) is 10 µm.

## Discussion

In this study, we investigated the mechanisms underlying the polar localization of cortical SOK (SOK) proteins using the SOK1 protein from *Arabidopsis thaliana* as a model. Through extensive mutagenesis, we probed the contributions of electrostatic interactions, specific amino acid sequences, and polymerization to the polar targeting of SOK1. We determined that polar targeting of SOK1 requires two components: the minimal polarity fragment and DIX-mediated polymerization. Our analysis of various mutants of the SOK1 polarity domain provided significant mechanistic insights. Most strikingly, we identified the precise minimal fragment required for polar localization of SOK1. This minimal SOK1 polarity motif, encoded by amino acids 213-237, is required and sufficient to target the chimeric protein to the inner apical edge of cortex cells when polymerized (e.g. via a DIX domain). To our knowledge, this is the first polar targeting sequence described in plant cells. Within this minimal polarity fragment, a combination of hydrophilic, positively charged, and hydrophobic residues polarly target the protein to the membrane.

*In silico* analysis of the minimal polarity fragment revealed that it encodes an amphipathic helix (AH), one face of which is hydrophobic, while the other face is positively charged. Such AHs are commonly found in membrane-binding proteins throughout the domains of life, from viruses to eukaryotes (Giménez-Andrés et al., 2018). The hydrophobic side of the helix is buried within membrane lipids, while the positively charged face interacts with the negative charges of the head groups. These amphipathic properties are essential for membrane association: removing positive residues in the polar face or removing hydrophobic residues in the apolar face of the AH of the polarity motif disrupts membrane targeting. An intriguing aspect of our findings is the potential interplay between the amphipathic helix and the post-translational modification of the Cysteine at its terminus. Previous studies of SOK1 showed that this residue is essential for membrane association (van Dop et al., 2020). More recently, we identified polarity-localized Protein Acyltransferase (PAT) enzymes that can target SOK1 to the PM in heterologous systems (Pukhovaya et al., 2026). Our data support a model where the AH is required for initial association of SOK1 to the PM, followed by PAT-dependent acylation. An attractive hypothesis is that the AH directly interacts with the membrane, yet with low affinity. This would bring SOK1 in close proximity to the PAT enzymes that subsequently stably anchor SOK1 to the PM. The effective reduction in diffusion from 3D to 2D may increase the probability of oligomerization, which lastly concentrates SOK1 protein in the cell edge.

There is a precedent for this interplay between an AH and PAT activity: a similar topology of an AH adjacent to a palmitoylated cysteine has been studied in several proteins. For example, the intracellular loop of the human sodium-calcium exchanger protein NCX1 contains an AH that regulates palmitoylation of adjacent cysteines. Mutations disrupting the AH also abolished palmitoylation. Moreover, fusion of this AH and the adjacent cysteine with YFP resulted in palmitoylation and membrane targeting of the fusion protein, demonstrating that this structure is a signal for S-acylation by DHHC-PAT enzymes (Plain et al., 2017). In a similar study, Lin et al. (Lin et al., 2015) identified an AH within the *Legionella* effector protein GobX. The association of this AH with the plasma membrane causes S-acylation of a cysteine within the AH and membrane association of the protein.

Whether this three-step model (1: AH-mediated PM association; 2: PAT-mediated acylation; 3 – DIX-mediated oligomerization) is sufficient to explain the native SOK patterns is an open question. It is possible that the AH in fact offers the interaction interface with the PAT enzyme, not the PM. Likewise, it is possible that either global or localized de-S-acylation may be required to maintain polarized SOK localization, analogous to the localized association and global dissociation of the yeast polar protein Cdc42 (Woods et al., 2016).

The identification of the minimal targeting region in SOK1 also allowed exploring the evolution of the polar targeting signals in the family. Through phylogenetic analysis and motif and sequence analysis, we found that SOK proteins fall into several ortholog groups, and that within angiosperms, the EuSOK-B group diversified into three distinct subgroups. A key feature shared by all SOKs is the presence of a Conserved Polarity Fragment (CPF) within their polarity domains (PDs), which drive their polarization. Synthesizing our findings, we propose a scenario for the evolutionary history of polar SOK proteins (**Fig. 6B**). Consistent with prior studies (Ramalho et al., 2021)(van Dop et al., 2020), our phylogenetic analysis shows that all extant SOKs originate from a single ancestral protein, AncSOK, at the base of land plants. This AncSOK gene evolved into BryoSOKs in bryophytes and into tracheophyte (vascular plant) SOKs. In vascular plants, a gene duplication event in early euphyllophytes gave rise to two major ortholog groups: EuSOK-A and EuSOK-B. These two groups have been broadly retained across different euphyllophyte lineages. A subsequent series of duplications in the EuSOK-B lineage resulted in three angiosperm-specific subgroups: EuSOK-B2, -B3, and -B4, which are retained in most angiosperms. Thus, while bryophytes possess at least one BryoSOK gene, ferns have at least two (EuSOK-A and EuSOK-B), and angiosperms typically have at least four SOK genes (one EuSOK-A and three EuSOK-Bs).

The evolutionary diversification of SOK proteins was accompanied by changes in their polar localization. Ancestral SOK proteins likely exhibited “inner” membrane localization, as inferred from the polarization patterns of modern BryoSOK, LycoSOK, EuSOK-A and fern EuSOK-B proteins. All tested polarity domains from these groups polarized to the inner membrane when expressed in *A. thaliana* root cells. This suggests that inner localization, or its equivalent in other species, was the ancestral state in AncSOK and was largely retained throughout SOK evolution. What exactly this “inner” domain corresponded to in the earliest land plants with vastly different morphology, remains an open question.

As EuSOK-Bs diversified into the angiosperm-specific subgroups their polar localization patterns shifted. For example, the inner basal localization of AtSOK2 (EuSOK-B2) and the uniform localization of AtSOK3 (EuSOK-B3) are newly evolved patterns. These shifts in localization are likely caused by the novel sequence motifs that evolved around the ancient CPF, possibly modifying interactions with additional cellular components. We note that this diversification appears restricted to the group that underwent gene duplication events. This suggests that duplications liberated SOK copies to diversify polarity patterns, whilst retaining at least one copy that kept the ancestral pattern. An open question is whether the *de novo* polarity patterns serve specialized functions, and if so, which function. Loss-of-function analysis will be required to answer this question.

Different SOK ortholog groups have evolved distinct patterns of charge and hydrophobicity within their polarity domains, reflecting underlying evolutionary trajectories. All EuSOK-A proteins encode amphipathic helices at the CPF. In AtSOK1, this amphipathic helix presumably enables the S-acylation of the conserved cysteine, which anchor the protein to the membrane. We argue that this mechanism likely extends to other EuSOK-A proteins as well. However, our phylogeny indicates that the amphipathic helix-dependent mechanism is an innovation specific to EuSOK-As, which is remarkable, since EuSOK-As seem to have retained the ancestral targeting localization properties of AncSOK. In contrast, EuSOK-B proteins and the earlier-diverging LycoSOK, retain an ancestral configuration of a hydrophobic CPF surrounded by positive charges. BryoSOKs differ even further, lacking a clear predicted helix and instead forming extended, unstructured loops with high positive charge around their CPF. Despite these differences, all studied MBMs drive polar localization, indicating that the conserved positive charge around the MBM is the primary driver of polarization across all SOK proteins. Interestingly, comparative proteomic analysis revealed that both Arabidopsis and Marchantia SOK proteins associate with PAT enzymes, and require the conserved Cysteine in the PD for polar membrane localization (Pukhovaya et al., 2026). Thus, while the initial targeting mechanisms to the PM (or PAT enzyme) may differ, there is deep conservation of the second and third steps in polar targeting.

In summary, our work identifies a minimal targeting region within SOK1, suggests a three-step polar targeting mechanism, and reveals the ancestral properties of SOK proteins and their evolutionary patterns. We anticipate that further analysis of the biochemical and biophysical mechanisms of SOK-PM interaction and loss-of-function analysis of the broader SOK family will be excellent starting points to understand general mechanisms underlying plant cell polarity.

## Materials and Methods

### Plants

*Arabidopsis thaliana* plants of the Columbia-0 ecotype were used for all experiments. For plate-based experiments, sterile seeds were grown on ½ MS + 1% sucrose medium and 1 % Agar (Daichin Agar, Duchefa). Seeds were stratified for 1-3 days at 4 °C, then placed vertically in a growth chamber in a 16 h / 8 h light / dark cycle at a temperature of 22°C. For propagation, 5-10 day old seedlings were transferred from plates to soil.

For generating transgenic plants, we used the floral dip method (Clough & Bent, 1998). In brief, *Agrobacterium tumefaciens* (strain GV3101 pSoup) carrying the construct of interest were grown to an OD of 0.5-0.8. Then, the cultures were diluted with an equal volume of 10% sucrose 0.1% Silwet L-77 solution. Plants with young bolts (corresponding to about 7-week-old plants) were then dipped in the resulting solution, allowed to recover for 1 day in a dark humid chamber, then replaced into the growth rooms.

Successfully transformed T_1_ plants were selected using an appropriate method depending on the construct used. Most of the constructs in this study use the FastGreen selection marker(Shimada et al., 2010). In this case, we selected green-fluorescent seeds under a stereomicroscope. For constructs with kanamycin resistance, T_1_ seeds were plated on plates supplemented with the 50 mg/L antibiotic. Surviving transformants were selected 10 days later.

*Marchantia polymorpha* Takaragaike-1 (Tak-1) was used as the wild-type accession for all Marchantia lines and experiments. Plants were cultured on ½ strength Gamborg B5 medium (Duchefa G0209) at 22°C under constant white light (40 μmol photons m⁻² s⁻¹).

Marchantia Tak-1 spores were transformed with the above constructs using *Agrobacterium tumefaciens* strain GV3101:pMp90, following a published protocol (Ishizaki et al., 2008). Transformants were selected on ½ strength B5 medium supplemented with 10 mg/L hygromycin and 100 μg/mL cefotaxime.

### Design and generation of expression constructs

#### Polarity domain constructs

All polarity domain constructs were expressed as mNeonGreen-tagged fusions using the RPS5a promoter, which drives expression in the root tips. To this end, we created a new expression vector. A C-terminal mNeonGreen tag was codon-optimized for plants and introduced into the previously published pPLV28 vector (De Rybel et al., 2011), which yielded the vector pNeonGreenKan. To avoid antibiotic-based selection and allow imaging of primary transformants, we replaced the resistance marker in this vector with a FastGreen selection marker. This new vector, pNeonGreenFG, was used as a backbone for further cloning rounds.

Constructs were generated as hDVL2-DIX DIX and mNeonGreen fusions, unless stated otherwise. For each construct, a synthetic DNA gBlock (Integrated DNA Technologies) encoding a variant of the SOK1 polarity domain, along with the hDvl2-DIX domain fragment, were used for cloning. These two regions were spaced with a five-amino-acid GGGSG linker. We assembled the constructs into the PstI site of pNeonGreenFG using HiFi cloning (New England Biolabs).

#### Polarity domain mutants

Targeted mutations were introduced into the polarity domain of SOK1 (amino acids 173-252). We substituted isolated amino acids with alanines (unless otherwise stated). To avoid disturbing the structure of the protein by introducing long stretches of hydrophobic alanines, we substituted longer fragments with flexible glycine-serine linkers. All mutant polarity domains were also expressed as hDvl2-DIX-polarity domain-mNeonGreen fusions as described above. Similarly, to manipulate the amphipathic properties of the SOK1 polarity motif, we introduced mutations in the SOK1 203-242 fragment, which we expressed as hDvl2-DIX-SOK1_203-242_-mNeonGreen fusions.

#### Polymerization domain constructs

The DIX domain in SOK1 was replaced with other polymerization domains and expressed as mNeonGreen-tagged fusions. We made three such constructs. The first replaces the SOK1 DIX domain with the DIX polymerization domain from the human DVL2 protein. Note that this construct corresponds to a C-terminal truncation of the previously published hDvl2-SOK1 construct (van Dop et al., 2020). The second construct uses a SAM polymerization domain (Sayou et al., 2016) form the *Arabidopsis thaliana* LFY protein (amino acids 37-158), which we obtained directly from cDNA. The third construct uses a PB1 polymerization domain from the *Marchantia polymorpha* ARF2 protein (Kato et al., 2020). Each construct was assembled in the HpaI restriction site of pNeonGreenK using HiFi cloning (new England Biolabs) and two inserts: a polymerization domain fragment and a fragment encoding the SOK1 polarity domain (amino acids 116-252).

#### Tandem oligomers

Two, four or six copies of the polarity domain were spaced with a flexible linker. The resulting tandem dimer, tetramer and hexamer were expressed with the same mNeonGreen tag but without any polymerization domain. To avoid issues resulting from repetitive DNA sequences, the individual units of the oligomers were encoded using different codons.

#### Cross-species polarity domain constructs

We expressed hDvl2-SOK polarity domain-mNeonGreen (mNG) fusion proteins under the *RPS5a* promoter as described above. The species and sequences are listed in **Table S2**.

An overview of all constructs used in this study can be found in Error! Reference source not found. **Table S1**.

### Microscopy and image analysis

Unless stated otherwise, we localized the various SOK proteins in roots of 5-day-old T_1_ or T_2_ seedlings, which represent independent insertion events of the transgene. To visualize cell outlines, cell walls were stained with 20 µg/ml Propidium iodide for 3 minutes. Then, the root tips were imaged on a Leica SP8 Dive multiphoton microscope using a Leica HC PL IRAPO 40x/1.10 water immersion objective. The settings of the microscope were as follows: excitation 940 nm, emission detection range: 500-560 nm (mNeonGreen) and 600-700 nm (propidium iodide) using HyD detectors with 10% Smart gain, scan format 1024 x 1024 pixels, 400 Hz. Images were acquired using the LAS X software, and processed in Fiji (Schindelin et al., 2012).

All images were only adjusted for contrast, brightness, and orientation, and these adjustments were applied uniformly to each image. Images with very low or no expression of the transgene were discarded from analysis. For each construct, one representative image is shown. The total number of independent lines imaged for each construct is indicated in each figure panel.

### Bioinformatic tools

The 3D structure of SOK1 (UniProt ID A0A178VZT8) was obtained from the AlphaFold Protein Structure Database (Stroe et al., 2021)(Varadi et al., 2022). Molecular graphics and analyses of the polarity domain structure were performed in UCSF ChimeraX version 1.7.1(Meng et al., 2023; Pettersen et al., 2021). HELIQUEST (Gautier et al., 2008) was used to perform amphipathicity analysis, hydrophobicity calculations and helical wheel projections. This analysis was performed using a 14 amino acid window. Prediction of peptide-membrane binding dynamics was performed using PMIpred (van Hilten et al., 2024).

### Analysis of SOK sequence data

We compiled a set of SOK sequences from various public databases (see Supplementary Methods for details on sequence sources). Most databases have Pfam/InterPro annotations, which allowed us to obtain proteins containing annotated SOSEKI DIX domains (Pfam ID: PF06136). Where no such annotations existed (*Metasequoia glyptostroboides, Lupinus angustifolius* and *Lycopodium clavatum*) we queried the genomes using BLASTP to identify sequences highly homologous to the *A. thaliana* SOK1 protein. Since genome sequences are available for few bryophyte species, we supplemented our dataset with additional bryophyte SOK sequences from the OneKP dataset (van Dop et al., 2020). We filtered out sequences that did not match a SOK DIX domain using PFAMScan (Madeira et al., 2024)(Mistry et al., 2020), as well as sequences with less than 200 amino acids (median sequence length before filtering: 427 amino acids). The resulting dataset contained 1086 sequences from 199 species.

Sequences were aligned using MAFFT L-Ins-2 (Katoh et al., 2017). This multiple sequence alignment (MSA) was used to build a phylogenetic tree using FastTree version 2.1.11 (Price et al., 2009). No trimming of the input sequences was performed. We used a JTT + gamma substitution model, which Smart Model Selection for PhyML (Lefort et al., 2017) showed as optimal. The resulting tree was used as a starting tree for GeneRax to infer a maximum likelihood species-tree-aware phylogeny. We ran GeneRax with the undated gene tree evolution model (UndatedDL) and optimized the tree topology using SPR moves with a maximum radius of 6.

We retrieved a species tree from OpenTree Taxonomy version 3.6 (https://zenodo.org/records/3937751). There were a few polytomies in the species tree. We resolved these polytomies for downstream use by grouping pairs of branches into artificial clades. Furthermore, we rerooted the species tree to reflect the monophyletic origin of bryophytes, which is currently the consensus topology of land plant phylogeny (Harris et al., 2020; Harris et al., 2022; Su et al., 2021). The species tree, corrected gene tree and the reconciled species-gene-tree showing duplications and losses can be found in the Supplementary materials. Gene trees are visualized and annotated using iTOL (Letunic and Bork, 2024). The reconciled gene family tree was visualized with ThirdKind version 3.6.8 (Penel et al., 2022).

### Data processing and visualization

Data was processed using R version 4.3.3 (Team, 2014) and RStudio version 2023.12.1+402 (Team, 2020). Graphics were generated using ggplot2 (Wickham, 2016). Images were processed using Fiji/ImageJ (Schindelin et al., 2012).

### Evolutionary conservation analysis

We used ConSurf (Ashkenazy et al., 2016; Berezin et al., 2004; Yariv et al., 2023) to calculate residue-level conservation of each ortholog group. For this, we selected the respective ortholog sequences of bryoSOKs, lycoSOKs, angiosperm sequences of euSOK-A and the three euSOK-B groups. Each sequence group was aligned using MAFFT. This alignment was used to calculate conservation scores using the ConSurf web service.

## Acknowledgements

The authors are grateful to Robertas Ursache and Niko Geldner for sharing plasmids, and for members of our team for helpful discussions.

## Data and resource availability

All primary data are available upon request from the corresponding author. Multiple sequence alignments are available at https://doi.org/10.6084/m9.figshare.28883615.v2. Sequences used are available at https://doi.org/10.6084/m9.figshare.28883615.v2. Phylogenetic trees are available at https://itol.embl.de/tree/193142201179291961725458366, https://itol.embl.de/tree/8610422102213921726438937 and https://doi.org/10.6084/m9.figshare.28883615.v2. Plasmid sequences are available at https://doi.org/10.6084/m9.figshare.34862703

## Supplementary Materials

**Figure S1:**
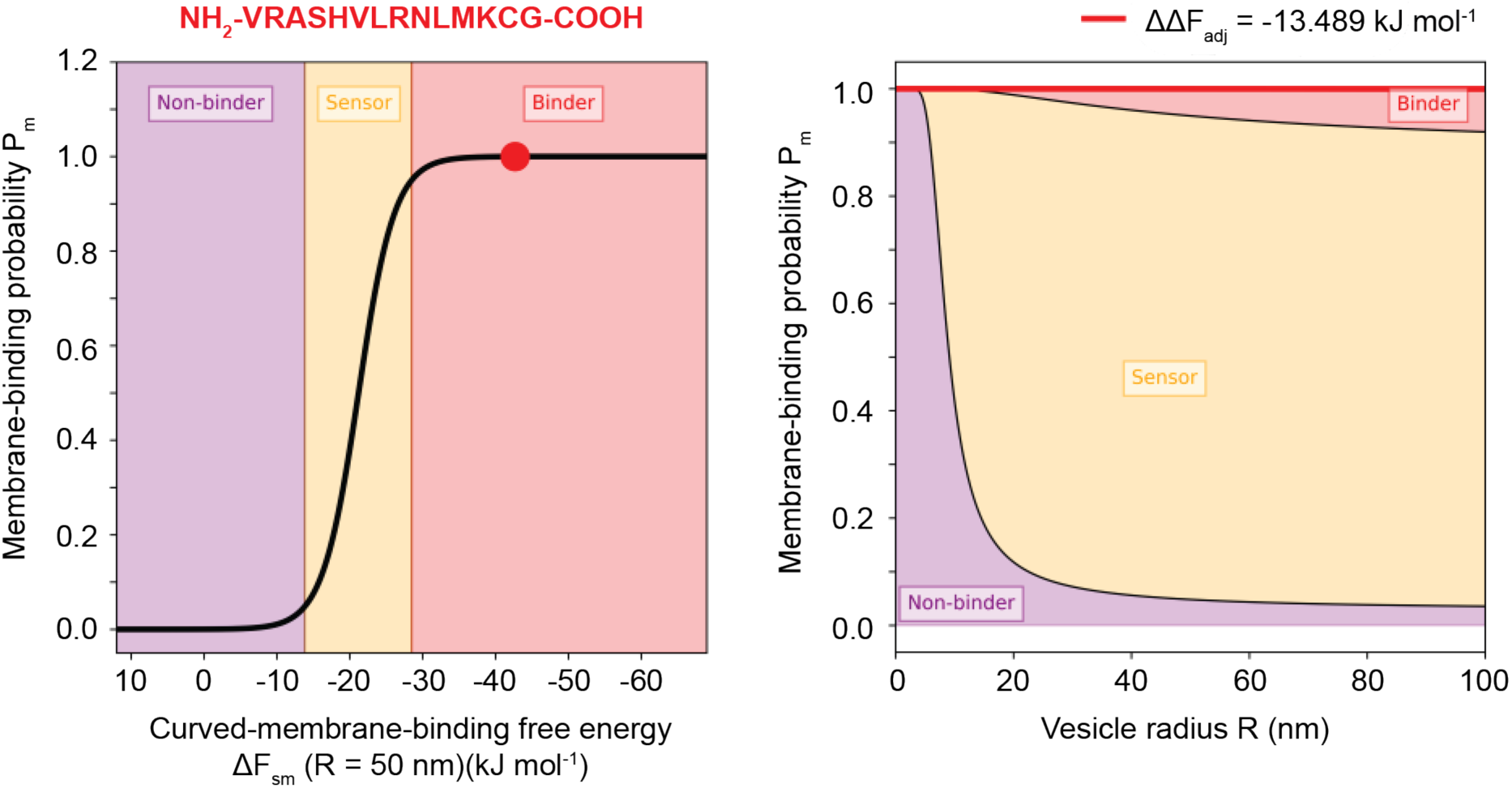
Predicted membrane binding properties of the SOK1 MPF. Plots depicting the membrane-binding probability of the SOK1-MPF as a function of membrane curvature (left) or vesicle radius (right). SOK1-MPF is indicated with a red dot (left) or line (right), relative to reference non-binder, sensor and binder peptides.

**Table S1:**
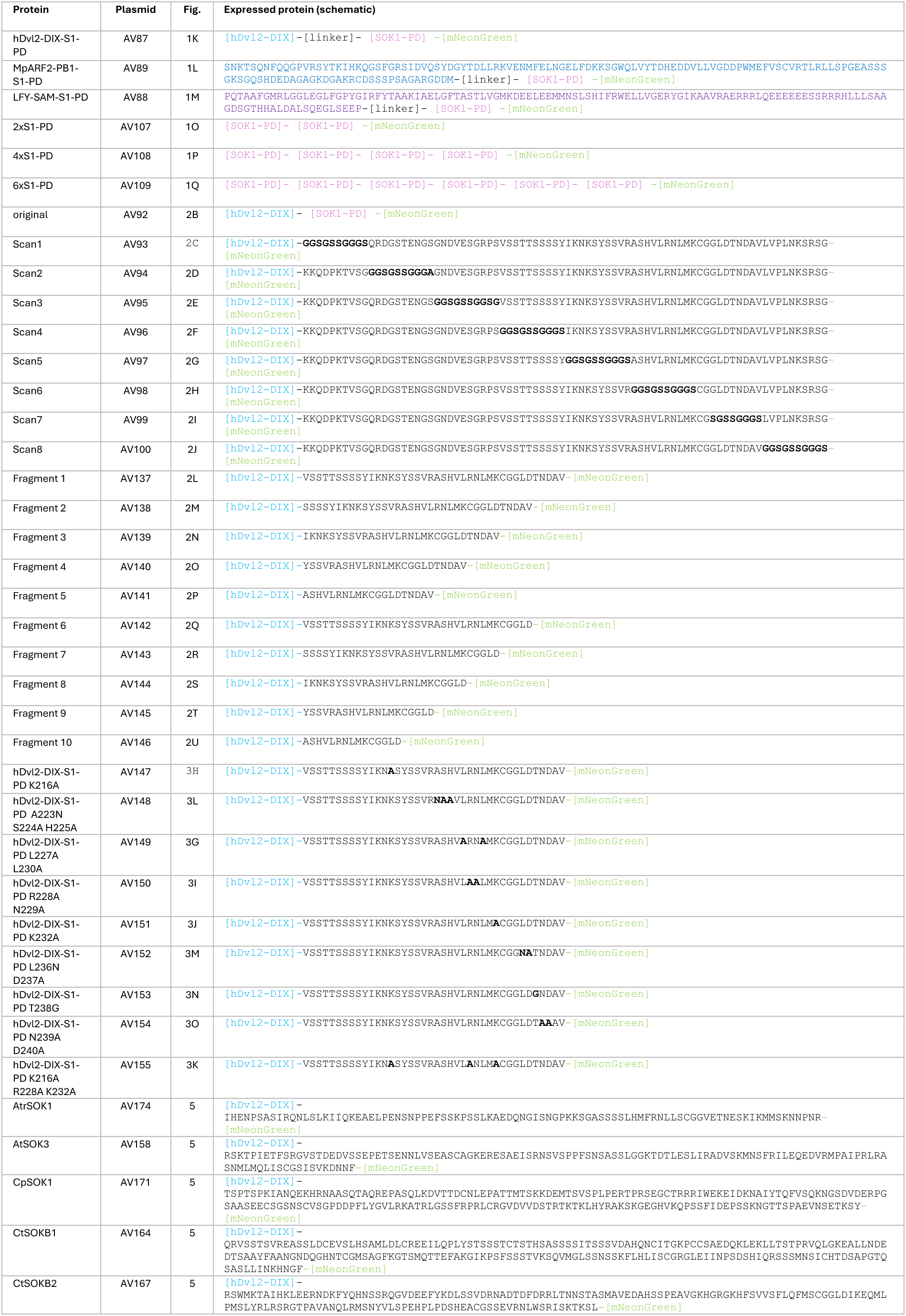
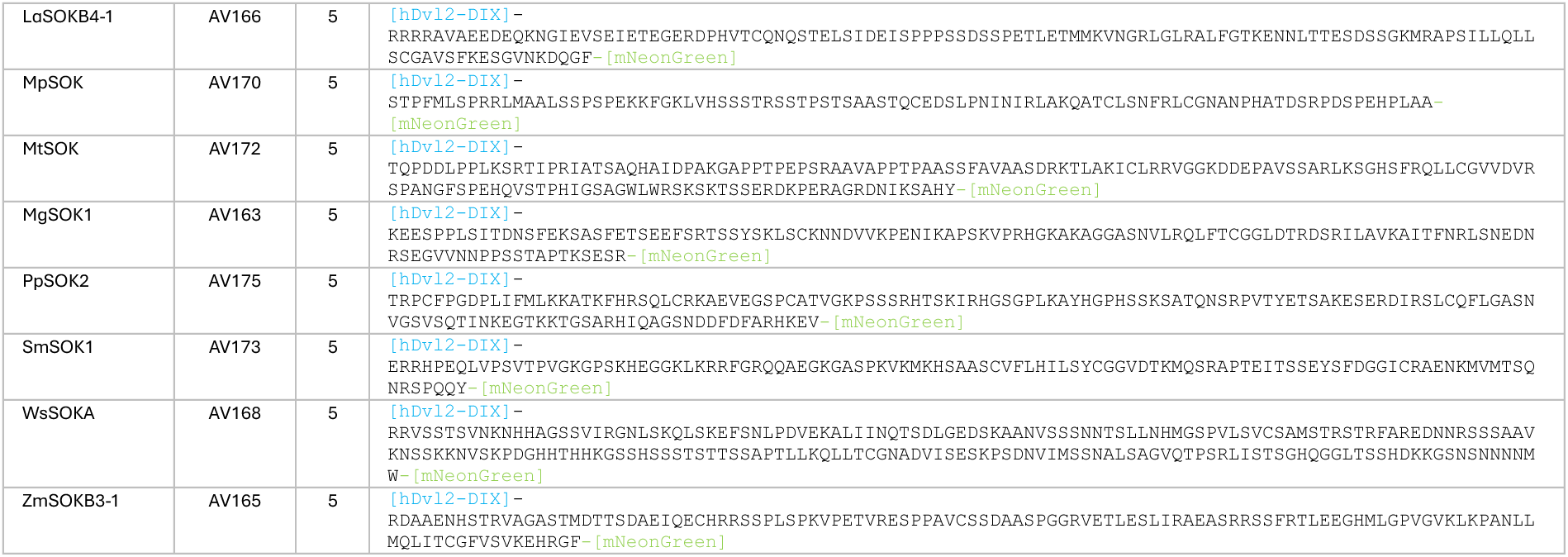
List of all constructs used in this study, with protein sequences.

| Protein | Plasmid | Fig. | Expressed protein (schematic) |
|---|---|---|---|
| hDvl2-DIX-S1-PD | AV87 | 1K | [hDvl2-DIX]-[linker]- [SOK1-PD] -[mNeonGreen] |
| MpARF2-PB1-S1-PD | AV89 | 1L | SNKTSQNFQQGPVRSYTKIHKQGSFGRSIDVQSYDGYTDLLRKVENMFELNGELFDKKSGWQLVYTDHEDDVLLVGDDPWMEFVSCVRTLRLLSPGEASSSGKSGQSHDEDAGAGKDGAKRCDSSSPSAGARGDDM-[linker]- [SOK1-PD] -[mNeonGreen] |
| LFY-SAM-S1-PD | AV88 | 1M | PQTAAFGMRLGGLEGLFGPYGIRFYTAAKIAELGFTASTLVGMKDEELEEMMNSLSHIFRWELLVGERYGIKAAVRAERRRLQEEEEEESSRRRHLLLSAAGDSGTHHALDALSQEGLSEEP-[linker]- [SOK1-PD] -[mNeonGreen] |
| 2xS1-PD | AV107 | 1O | [SOK1-PD]- [SOK1-PD] -[mNeonGreen] |
| 4xS1-PD | AV108 | 1P | [SOK1-PD]- [SOK1-PD]- [SOK1-PD]- [SOK1-PD] -[mNeonGreen] |
| 6xS1-PD | AV109 | 1Q | [SOK1-PD]- [SOK1-PD]- [SOK1-PD]- [SOK1-PD]- [SOK1-PD]- [SOK1-PD] -[mNeonGreen] |
| original | AV92 | 2B | [hDvl2-DIX]- [SOK1-PD] -[mNeonGreen] |
| Scan1 | AV93 | 2C | [hDvl2-DIX]-**GGSGSSGGGS**QRDGSTENGSGNDVESGRPSVSSTTSSSSYIKNKSYSSVRASHVLRNLMKCGGLDTNDAVLVPLNKSRSG-[mNeonGreen] |
| Scan2 | AV94 | 2D | [hDvl2-DIX]-KKQDPKTVSG**GGSGSSGGGA**GNDVESGRPSVSSTTSSSSYIKNKSYSSVRASHVLRNLMKCGGLDTNDAVLVPLNKSRSG-[mNeonGreen] |
| Scan3 | AV95 | 2E | [hDvl2-DIX]-KKQDPKTVSGQRDGSTENGS**GGSGSSGGSG**VSSTTSSSSYIKNKSYSSVRASHVLRNLMKCGGLDTNDAVLVPLNKSRSG-[mNeonGreen] |
| Scan4 | AV96 | 2F | [hDvl2-DIX]-KKQDPKTVSGQRDGSTENGSGNDVESGRPS**GGSGSSGGGS**IKNKSYSSVRASHVLRNLMKCGGLDTNDAVLVPLNKSRSG-[mNeonGreen] |
| Scan5 | AV97 | 2G | [hDvl2-DIX]-KKQDPKTVSGQRDGSTENGSGNDVESGRPSVSSTTSSSSY**GGSGSSGGGS**ASHVLRNLMKCGGLDTNDAVLVPLNKSRSG-[mNeonGreen] |
| Scan6 | AV98 | 2H | [hDvl2-DIX]-KKQDPKTVSGQRDGSTENGSGNDVESGRPSVSSTTSSSSYIKNKSYSSVR**GGSGSSGGGS**CGGLDTNDAVLVPLNKSRSG-[mNeonGreen] |
| Scan7 | AV99 | 2I | [hDvl2-DIX]-KKQDPKTVSGQRDGSTENGSGNDVESGRPSVSSTTSSSSYIKNKSYSSVRASHVLRNLMKCG**SGSSGGGS**LVPLNKSRSG-[mNeonGreen] |
| Scan8 | AV100 | 2J | [hDvl2-DIX]-KKQDPKTVSGQRDGSTENGSGNDVESGRPSVSSTTSSSSYIKNKSYSSVRASHVLRNLMKCGGLDTNDAV**GGSGSSGGGS**-[mNeonGreen] |
| Fragment 1 | AV137 | 2L | [hDvl2-DIX]-VSSTTSSSSYIKNKSYSSVRASHVLRNLMKCGGLDTNDAV-[mNeonGreen] |
| Fragment 2 | AV138 | 2M | [hDvl2-DIX]-SSSSYIKNKSYSSVRASHVLRNLMKCGGLDTNDAV-[mNeonGreen] |
| Fragment 3 | AV139 | 2N | [hDvl2-DIX]-IKNKSYSSVRASHVLRNLMKCGGLDTNDAV-[mNeonGreen] |
| Fragment 4 | AV140 | 2O | [hDvl2-DIX]-YSSVRASHVLRNLMKCGGLDTNDAV-[mNeonGreen] |
| Fragment 5 | AV141 | 2P | [hDvl2-DIX]-ASHVLRNLMKCGGLDTNDAV-[mNeonGreen] |
| Fragment 6 | AV142 | 2Q | [hDvl2-DIX]-VSSTTSSSSYIKNKSYSSVRASHVLRNLMKCGGLD-[mNeonGreen] |
| Fragment 7 | AV143 | 2R | [hDvl2-DIX]-SSSSYIKNKSYSSVRASHVLRNLMKCGGLD-[mNeonGreen] |
| Fragment 8 | AV144 | 2S | [hDvl2-DIX]-IKNKSYSSVRASHVLRNLMKCGGLD-[mNeonGreen] |
| Fragment 9 | AV145 | 2T | [hDvl2-DIX]-YSSVRASHVLRNLMKCGGLD-[mNeonGreen] |
| Fragment 10 | AV146 | 2U | [hDvl2-DIX]-ASHVLRNLMKCGGLD-[mNeonGreen] |
| hDvl2-DIX-S1-PD K216A | AV147 | 3H | [hDvl2-DIX]-VSSTTSSSSYIKN**A**SYSSVRASHVLRNLMKCGGLDTNDAV-[mNeonGreen] |
| hDvl2-DIX-S1-PD A223N S224A H225A | AV148 | 3L | [hDvl2-DIX]-VSSTTSSSSYIKNKSYSSVR**NAA**VLRNLMKCGGLDTNDAV-[mNeonGreen] |
| hDvl2-DIX-S1-PD L227A L230A | AV149 | 3G | [hDvl2-DIX]-VSSTTSSSSYIKNKSYSSVRASHV**A**RN**A**MKCGGLDTNDAV-[mNeonGreen] |
| hDvl2-DIX-S1-PD R228A N229A | AV150 | 3I | [hDvl2-DIX]-VSSTTSSSSYIKNKSYSSVRASHVL**AA**LMKCGGLDTNDAV-[mNeonGreen] |
| hDvl2-DIX-S1-PD K232A | AV151 | 3J | [hDvl2-DIX]-VSSTTSSSSYIKNKSYSSVRASHVLRNLM**A**CGGLDTNDAV-[mNeonGreen] |
| hDvl2-DIX-S1-PD L236N D237A | AV152 | 3M | [hDvl2-DIX]-VSSTTSSSSYIKNKSYSSVRASHVLRNLMKCGG**NA**TNDAV-[mNeonGreen] |
| hDvl2-DIX-S1-PD T238G | AV153 | 3N | [hDvl2-DIX]-VSSTTSSSSYIKNKSYSSVRASHVLRNLMKCGGLD**G**NDAV-[mNeonGreen] |
| hDvl2-DIX-S1-PD N239A D240A | AV154 | 3O | [hDvl2-DIX]-VSSTTSSSSYIKNKSYSSVRASHVLRNLMKCGGLDT**AA**AV-[mNeonGreen] |
| hDvl2-DIX-S1-PD K216A R228A K232A | AV155 | 3K | [hDvl2-DIX]-VSSTTSSSSYIKN**A**SYSSVRASHVL**A**NLM**A**CGGLDTNDAV-[mNeonGreen] |
| AtrSOK1 | AV174 | 5 | [hDvl2-DIX]-IHENPSASIRQNLSLKIIQKEAELPENSNPPEFSSKPSSLKAEDQNGISNGPKKSGASSSSLHMFRNLLSCGGVETNESKIKMMSKNNPNR-[mNeonGreen] |
| AtSOK3 | AV158 | 5 | [hDvl2-DIX]-RSKTPIETFSRGVSTDEDVSSEPETSENNLVSEASCAGKERESAEISRNSVSPPFSNSASSLGGKTDTLESLIRADVSKMNSFRILEQEDVRMPAIPRLRASNMLMQLISCGSISVKDNNF-[mNeonGreen] |
| CpSOK1 | AV171 | 5 | [hDvl2-DIX]-TSPTSPKIANQEKHRNAASQTAQREPASQLKDVTTDCNLEPATTMTSKKDEMTSVSPLPERTPRSEGCTRRRIWEKEIDKNAIYTQFVSQKNGSDVDERPGSAASEECSGSNSCVSGPDDPFLYGVLRKATRLGSSFRPRLCRGVDVVDSTRTKTKLHYRAKSKGEGHVKQPSSFIDEPSSKNGTTSPAEVNSETKSY-[mNeonGreen] |
| CtSOKB1 | AV164 | 5 | [hDvl2-DIX]-QRVSSTSVREASSLDCEVSLHSAMLDLCREEILQPLYSTSSSTCTSTHSASSSSITSSSVDAHQNCITGKPCCSAEDQKLEKLLTSTPRVQLGKEALLNDEDTSAAYFAANGNDQGHNTCGMSAGFKGTSMQTTEFAKGIKPSFSSSTVKSQVMGLSSNSSKFLHLISCGRGLEIINPSDSHIQRSSSMNSICHTDSAPGTQSASLLINKHNGF-[mNeonGreen] |
| CtSOKB2 | AV167 | 5 | [hDvl2-DIX]-RSWMKTAIHKLEERNDKFYQHNSSRQGVDEEFYKDLSSVDRNADTDFDRRLTNNSTASMAVEDAHSSPEAVGKHGRGKHFSVVSFLQFMSCGGLDIKEQMLPMSLYRLRSRGTPAVANQLRMSNYVLSPEHPLPDSHEACGSSEVRNLWSRISKTKSL-[mNeonGreen] |
| LaSOKB4-1 | AV166 | 5 | [hDvl2-DIX]-RRRRAVAEEDEQKNGIEVSEIETEGERDPHVTCQNQSTELSIDEISPPPSSDSSPETLETMMKVNGRLGLRALFGTKENNLTTESDSSGKMRAPSILLQLLSCGAVSFKESGVNKDQGF-[mNeonGreen] |
| MpSOK | AV170 | 5 | [hDvl2-DIX]-STPFMLSPRRLMAALSSPSPEKKFGKLVHSSSTRSSTPSTSAASTQCEDSLPNINIRLAKQATCLSNFRLCGNANPHATDSRPDSPEHPLAA-[mNeonGreen] |
| MtSOK | AV172 | 5 | [hDvl2-DIX]-TQPDDLPPLKSRTIPRIATSAQHAIDPAKGAPPTPEPSRAAVAPPTPAASSFAVAASDRKTLAKICLRRVGGKDDEPAVSSARLKSGHSFRQLLCGVVDVRSPANGFSPEHQVSTPHIGSAGWLWRSKSKTSSERDKPERAGRDNIKSAHY-[mNeonGreen] |
| MgSOK1 | AV163 | 5 | [hDvl2-DIX]-KEESPPLSITDNSFEKSASFETSEEFSRTSSYSKLSCKNNDVVKPENIKAPSKVPRHGKAKAGGASNVLRQLFTCGGLDTRDSRILAVKAITFNRLSNEDNRSEGVVNNPPSSTAPTKSESR-[mNeonGreen] |
| PpSOK2 | AV175 | 5 | [hDvl2-DIX]-TRPCFPGDPLIFMLKKATKFHRSQLCRKAEVEGSPCATVGKPSSSRHTSKIRHGSGPLKAYHGPHSSKSATQNSRPVTYETSAKESERDIRSLCQFLGASNVGSVSQTINKEGTKKTGSARHIQAGSNDDFDFARHKEV-[mNeonGreen] |
| SmSOK1 | AV173 | 5 | [hDvl2-DIX]-ERRHPEQLVPSVTPVGKGPSKHEGGKLKRRFGRQQAEGKGASPKVKMKHSAASCVFLHILSYCGGVDTKMQSRAPTEITSSEYSFDGGICRAENKMVMTSQNRSPQQY-[mNeonGreen] |
| WsSOKA | AV168 | 5 | [hDvl2-DIX]-RRVSSSTSVNKNHHAGSSVIRGNLSKQLSKEFSNLPDVEKALIINQTSDLGEDSKAANVSSSNNTSLLNHMGSPVLSVCSAMSTRSTRFAREDNNRSSSAAVKNSSKKNVSKPDGHHTHHKGSSHSSSTSTTSSAPTLLKQLLTCGNADVISESKPSDNVIMSSNALSAGVQTPSRLISTSGHQGGLTSSHDKKGSNSNNNNMW-[mNeonGreen] |
| ZmSOKB3-1 | AV165 | 5 | [hDvl2-DIX]-RDAAENHSTRVAGASTMDTTSDAEIQECHRRSSPLSPKVPETVRESPPAVCSSDAASPGGRVETLESLIRAEASRSSFRTLEEGHMLGPVGVKLKPANLLMQLITCGFVSVKEHRGF-[mNeonGreen] |

**Table S2:** Polarity domains used in this study. Shown are abbreviated sample names used in the main text (for bioinformatic analysis and microscopy of hDvl2-polarity domain-mNG fusions), corresponding species and gene or sequence ID, as well as the amino acids corresponding to the polarity domain. In some cases, we made more than one construct per gene, and the respective polarity fragments are given in the table.

| **Protein** | **Type** | **Species** | **Gene or Sequence ID** | **Region** |
|---|---|---|---|---|
| AtrSOK1 | EuSOKA | *Amborella trichopoda* | AmTrH1.03G055500 | 192-282 |
| AtSOK1 | EuSOKA | *Arabidopsis thaliana* | AT1G05577 | 173-252 |
| AtSOK3 | EuSOKB3 | *Arabidopsis thaliana* | AT2G28150 | 232-352 |
| CpSOK1 | BryoSOK | *Ceratodon purpureus* | CepurR40.11G122000 | 257-454 |
| CtSOKB1 | EuSOKB | *Ceratopteris thalictroides* | LM_PIVW_2013750_Ceratopteris_thalictroides_g.430 | 319-532 |
| CtSOKB2 | EuSOKB | *Ceratopteris thalictroides* | LM_PIVW_2008561_Ceratopteris_thalictroides_g.428 | 391-548 |
| LaSOKB4-1 | EuSOKB4 | *Lupinus angustifolius* | Lupan_006625 | 193-311 |
| MpSOK | BryoSOK | *Marchantia polymorpha* | Mapoly0032s0007 | 423-515 |
| MtSOK | BryoSOK | *Megaceros tosanus* | HW_UCRN_2009276_Megaceros_tosanus_g.336 | 241-391 |
| MgSOK1 | EuSOKA | *Metasequoia glyptostroboides* | Mgl0103920 | 239-360 |
| PpSOK2 | BryoSOK | *Physcomitrium patens* | Pp3c2_20380 | 310-448 |
| SmSOK1 | LycoSOK | *Selaginella moellendorffii* | 412562 | 293-400 |
| WsSOKA | EuSOKA | *Woodsia scopulina* | LM_YJJY_2005726_Woodsia_scopulina_g.459 | 391-593 |
| ZmSOKB3-1 | EuSOKB3 | *Zea mays* | Zm00001eb039020 | 292-409 |

